# Plasmid-encoded anatoxin biosynthesis in benthic cyanobacteria

**DOI:** 10.64898/2026.06.26.734624

**Authors:** Florian Gibis, Dominik Rieth, Juergen Geist, Alexandra Schoenle, Franziska Bauer, Charlotte Schampera, Ann-Marie Waldvogel

## Abstract

Anatoxin-producing cyanobacteria pose growing ecological and public health risks, yet the genomic organization of anatoxin biosynthesis remains poorly resolved. Using complete genome assembly of a benthic cyanobacterium of the genus *Microcoleus*, we show that the entire anatoxin biosynthetic gene cluster is encoded on a small circular plasmid rather than the chromosome. This unexpected localization suggests enhanced mobility of anatoxin biosynthesis and has implications for toxin evolution, release, and environmental surveillance.

## Main text

Cyanobacteria produce a diverse array of secondary metabolites, including some of the most potent natural neurotoxins known (Pearson et al. 2016). Among these, anatoxins pose an increasing global ecological and public health concern due to their rapid mode of action and association with cyanobacterial proliferations in freshwater ecosystems (Huisman et al. 2018). Although the biosynthetic gene cluster responsible for anatoxin production has been characterized in several planktonic and benthic cyanobacteria (Rantala-Ylinen et al. 2011; Méjean et al. 2009), it is generally assumed to be chromosomally encoded. Whether alternative genomic architectures contribute to the evolution, mobility, or environmental distribution of anatoxin biosynthesis remains unclear.

Here, we isolated and cultivated a benthic cyanobacterial strain (*Microcoleus* sp. MC67) associated with a fatal dog intoxication event in a German freshwater reservoir (Bauer et al. 2020). Using long-read sequencing, we generated a complete, circularized genome assembly comprising two replicons: a 5.61 Mb chromosome with 540 × coverage and a 49 kb circular extrachromosomal element with 108 × coverage. Assembly quality metrics, read back-mapping, and completeness assessments confirmed a high-quality, contamination-free genome, enabling unambiguous resolution of repetitive regions and secondary replicons (Supplementary Note 2). Phylogenomic analysis placed strain MC67 within the *Microcoleus* clade, closely related to *Microcoleus bour-rellyi* FEM_GT703, consistent with its filamentous morphotype (Supplementary Fig. 1).

Unexpectedly, genome mining revealed that the entire anatoxin biosynthetic gene cluster was located on the 49 kb circular element rather than on the chromosome (Fig. 1a). AntiSMASH analysis identified a complete dihydroanatoxin-a (dhATX)–like gene cluster spanning genes *anaA* through *anaK*, with gene content and synteny closely resembling previously characterized dhATX–producing clusters from *Microcoleus* and *Cylindrospermum* strains (Fig. 1b). Comparative analysis showed high amino-acid similarity (95–100% for core biosynthetic genes) to the dhATX cluster of *Microcoleus anatoxicus* PTRS3 (Fig. 1c). The cluster lacked *anaH*, a transposase-associated gene that is variably present among anatoxin-producing cyanobacteria (Brown et al. 2016), but retained all genes required for anatoxin-a (ATX) and dhATX biosynthesis (Méjean et al. 2016)(Supplementary Note 4).

**Fig. 1.**
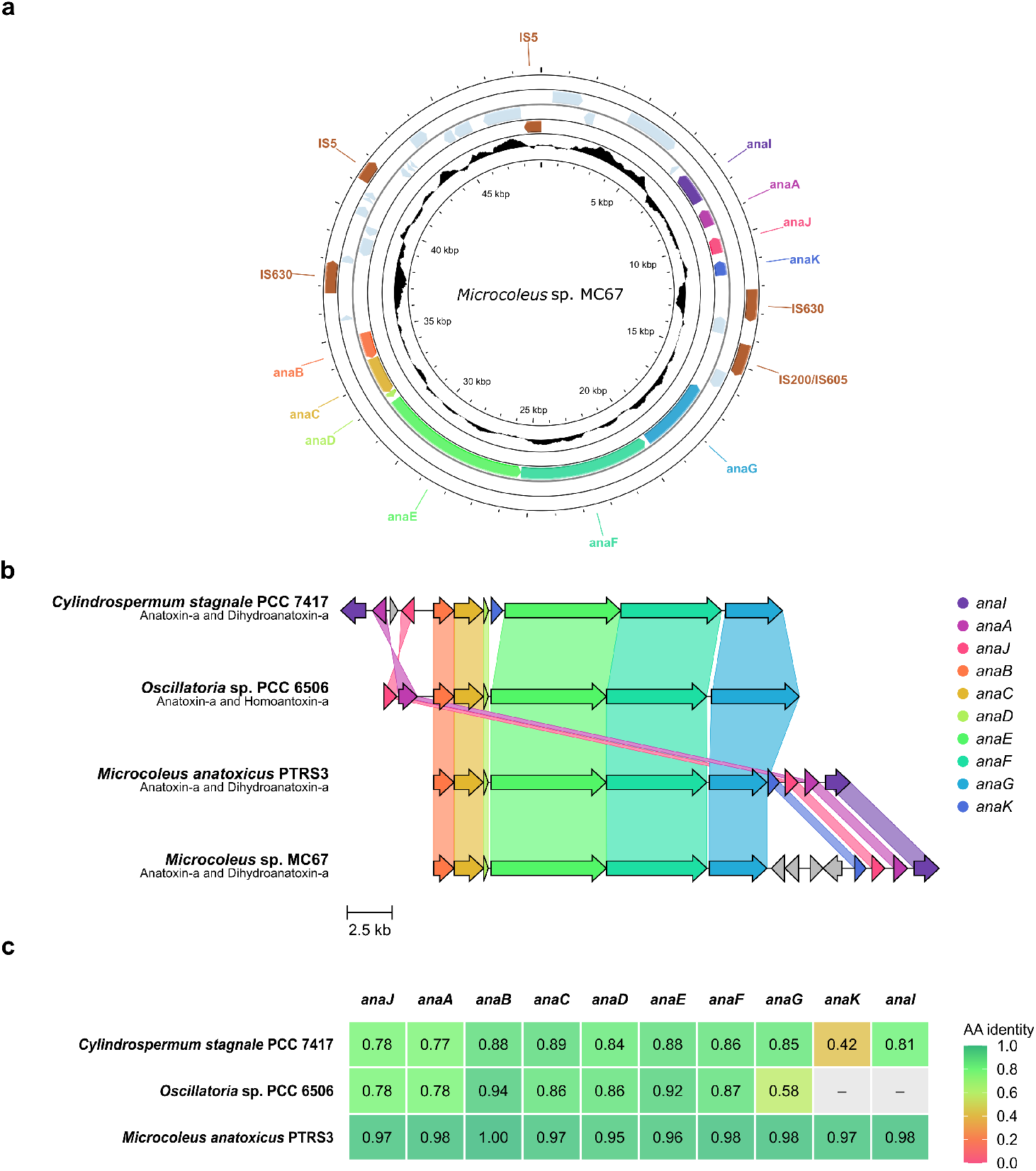
**a**: *Microcoleus* sp. MC67, containing the dihydroanatoxin-a cluster flanked by several insertion sequences from different families. Transparent features represent coding DNA sequences (CDS), GC content is shown on the innermost circle. IS denotes insertion sequence elements located on the plasmid. **b**: Comparison of the antiSMASH anatoxin-a reference cluster from *Oscillatoria* sp. PCC 6506, the dihydroanatoxin-a producing clusters of *Cylindrospermum stagnale* PCC 7417 and *Microcoleus anatoxicus* PTRS3, and the plasmid-like contig of *Microcoleus* sp. MC67. **c**: Amino acid identity of Figure **b** compared to *Microcoleus* sp. MC67, calculated by clinker.

Multiple lines of evidence supported the extrachromosomal nature of the anatoxin gene cluster (Supplementary Note 3). The 49 kb contig was circular, exhibited distinct coverage from the chromosome (Antipov et al. 2016), and was classified as plasmid-like by PlasFlow with a probability of 70%. Although no known replication or mobilization genes were detected by MOB-typer, the element contained numerous insertion sequence (IS) elements flanking the biosynthetic genes (Fig. 1a), indicating a dynamic genomic environment. Such features suggest that the anatoxin cluster resides on a mobile or formerly mobile genetic element rather than representing a chromosomal rearrangement or assembly artifact.

Chemical analyses provided functional support for the plasmid-encoded biosynthetic capacity. Among all targeted cyanotoxins, LC-MS/MS only detected ATX and dhATX; dhATX dominated strongly (192 to 2,420 *µ*g kg^*−*1^ sample wet mass), whereas ATX occurred at markedly lower concentrations (0.22 to 12.16 *µ*g kg^*−*1^ sample wet mass). The presence of *anaK*, encoding a putative F_420_-dependent oxidoreductase previously implicated in dhATX formation (Kust et al. 2020), is consistent with the observed toxin profile.

Phylogenomic analysis placed strain MC67 within the *Microcoleus* clade and showed that anatoxin production is distributed across multiple cyanobacterial lineages (Fig. 2), consistent with a dynamic evolutionary history of anatoxin biosynthesis. The concordance between genomic potential and toxin production in MC67, together with its phylogenetic context(Supplementary Note 1), supports the view that anatoxin biosynthetic gene clusters are evolutionarily flexible traits that can be maintained or acquired across divergent lineages (Popin et al. 2021).

**Fig. 2.**
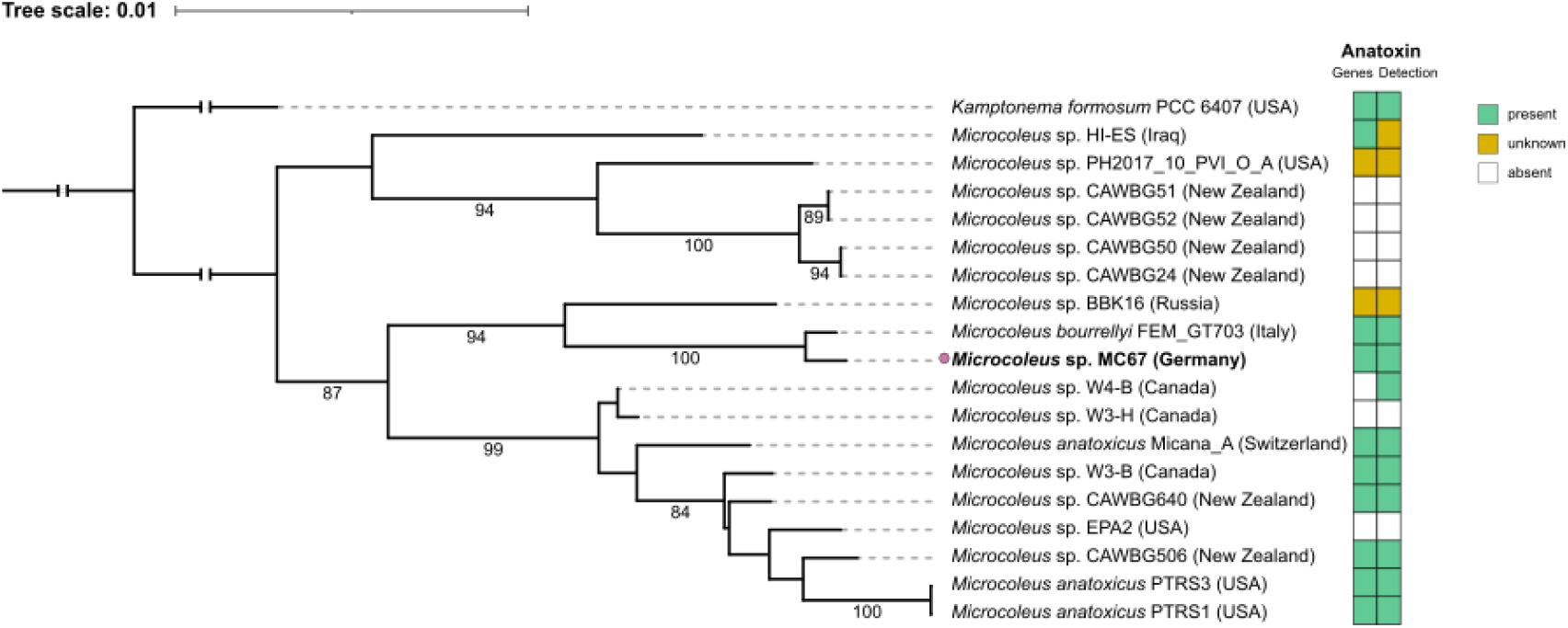
Maximum-likelihood (ML) phylogeny inferred from the concatenated GTDB `bac120` amino-acid marker alignment of 19 genome assemblies. The tree was midpoint-rooted, with *Kamptonema formosum* PCC 6407 included as outgroup. Ultrafast bootstrap support values ≥ 70% are displayed on branches. Scale bar (top) represents 0.01 expected substitutions per site in the ML analysis. Tip labels indicate strain name, country, and sampling locality; taxonomic names follow GTDB taxonomy. The adjacent two-column heatmap summarizes toxin-status information, with the first column indicating the presence of anatoxin biosynthesis genes and the second column indicating anatoxin detection. Green indicates presence, white indicates absence, and yellow indicates unknown or unavailable information.

Plasmid localization of a complete and functional anatoxin biosynthetic gene cluster represents a striking departure from the canonical chromosomal organization reported for cyanobacterial toxin genes. While large cyanobacterial plasmids (> 100 kb) have occasionally been shown to harbor biosynthetic gene clusters (Popin et al. 2021), the occurrence of an intact anatoxin cluster on a small (∼49 kb) plasmid-like element has not, to the best of our knowledge, been shown previously. The presence of multiple IS elements flanking key biosynthetic genes further suggests that recombination and horizontal gene transfer may have contributed to the assembly or dissemination of this cluster (Entfellner et al. 2022).

These findings have important implications for the evolution and environmental spread of cyanobacterial neurotoxins. Plasmid-encoded toxin biosynthesis could facilitate rapid horizontal transfer between species and lineages, potentially accelerating the emergence of toxic phenotypes in benthic cyanobacterial communities (Popin et al. 2021). From a monitoring perspective, extrachromosomal localization of toxin genes may complicate genomic surveillance strategies that rely on chromosomal markers alone. More broadly, our results highlight the need to consider plasmids as active contributors to the ecology and evolution of cyanobacterial secondary metabolism, particularly in benthic systems that remain underrepresented in genomic and toxicological studies (Wood et al. 2020).

## Methods

### Strain origin and cultivation

Strain MC67 was isolated from benthic cyanobacterial aggregates collected from littoral sediment of reservoir Mandichosee (Germany; 48^*°*^15^*′*^25.1^*′′*^N, 10^*°*^56^*′*^15.3^*′′*^E) in July 2021. Cultures were maintained in Z8 medium (Rippka 1988) at 15 °C under a 16 h light/8 h dark photoperiod (20 µmol m^*−*2^ s^*−*1^).

### DNA extraction and sequencing

High-molecular-weight DNA was obtained using bead-beating, enzymatic lysis, and phenol–chloroform extraction. Detailed extraction procedures are provided in the Supplementary Information (Supplementary Note 5). DNA quantity was measured using a Qubit fluorometer and fragment length was assessed using a TapeStation system. Libraries were prepared using the Oxford Nanopore Technologies ligation sequencing kit (v14) and sequenced on a MinION Mk1D platform, yielding approximately 5 Gb of long-read data.

### Genome assembly and annotation

Reads were basecalled with Dorado, filtered using Filtlong (Boostrom et al. 2022), and assembled with Flye (Kolmogorov et al. 2019). Assemblies were polished with NextPolish (Hu et al. 2020) using Nanopore and Illumina reads. Read mapping was performed with minimap2 (Li 2018) and bwa-mem (Li and Durbin 2009) to assess assembly consistency and coverage. Genome completeness and contamination were evaluated using BUSCO (Simão et al. 2015), CheckM2 (Parks et al. 2015), and QUAST (Mikheenko et al. 2018). Genome annotation was performed with Bakta (Schwengers et al. 2021). Biosynthetic gene clusters were identified using antiSMASH (Medema et al. 2011). Gene cluster comparisons and genome visualization were performed using clinker (Gilchrist and Chooi 2021) and Proksee (Grant et al. 2023). Reference sequences used for anatoxin biosynthetic gene cluster comparison are listed in Supplementary Table 2. Insertion sequences were detected using ISEScan (Xie and Tang 2017), and plasmid characteristics were inferred using PlasFlow (Krawczyk et al. 2018) and MOB-typer (Robertson and Nash 2018). Program parameters are provided in the Supplementary Material (Supplementary Note 6).

### Phylogenetic analysis

Phylogenetic placement was performed using GTDB-Tk (Chaumeil et al. 2022) based on the bac120 marker gene set and publicly available *Microcoleus* genomes. Maximum-likelihood phylogenies were inferred with IQ-TREE (Minh et al. 2020) using ModelFinder (Kalyaanamoorthy et al. 2017) and 1,000 ultrafast bootstrap replicates (Hoang et al. 2018). Phylogenies were visualized with iTOL (Letunic and Bork 2024). Pairwise average nucleotide identity (ANI) was calculated using FastANI (Jain et al. 2018). The accession numbers of all genomes included in the phylogenetic tree are listed in Supplementary Table 1.

### Toxin analysis

Cyanobacterial samples were acidified to stabilize the toxins and subsequently extracted using repeated freeze–thaw cycles combined with ultrasonication. Extracts were filtered (PVDF, 0.45 µm) prior to analysis. Toxins were quantified by LC-MS/MS following established protocols (Fastner et al. 2023), inlcuding the detection and quantification of anatoxin-a (ATX) and dihydroanatoxin-a (dhATX) by multiple reaction monitoring (MRM).

## Data availablity

The assembly was deposited in the European Nucleotide Archive (ENA); accession number and taxonomic ID assignment are pending.

## Data visualization

Figures were generated using R (R Core Team 2025), Proksee (Grant et al. 2023), and clinker (Gilchrist and Chooi 2021), with final layouts prepared in Inkscape.

Detailed experimental protocols and extended methodological descriptions are provided in the Supplementary Information.

## Acknowledgements

The authors sincerely thank David Konkol (Umweltbundesamt, Berlin) for his skilled technical assistance in the laboratory and Michael Wurm (Technical University, Munich) for the isolation of the strain. We also gratefully acknowledge the computational and data resources as well as the support provided by the Leibniz Supercomputing Centre (www.lrz.de).

## Author contributions

Conceptualization: F.G., D.R., J.G. and A.M.W.; Methodology: F.G., D.R, A.S. and C.S.; Analysis: F.G. and D.R.; Visualization: F.G. and D.R.; Writing: A.M.W., F.G. and D.R.; Funding: J.G., A.M.W; Supervision: J.G.; A.M.W.; Reviewing and editing the manuscript: all authors

## Funding

The research project (CyanoBenToxII) is funded by the Bavarian State Ministry of Health, Care, and Prevention (StMGP), and the DIVATOX-2 project funded by the German Federal Ministry of Education and Research under funding code 16LW0542K. The funders had no role in study design; in the collection, analysis, or interpretation of data; in the writing of the manuscript; or in the decision to submit the article for publication.

## Competing interests

The authors declare no competing interests.

## Additional information

**Supplementary information** The online version contains supplementary material available at XXX.

## Notes

### Competing Interest Statement

The authors have declared no competing interest.

