## Supplementary materials for "Plasmid-encoded anatoxin biosynthesis in benthic cyanobacteria"

#### Supplementary Figure 1

##### Macroscopic and microscopic morphology of *Microcoleus* sp. MC67.

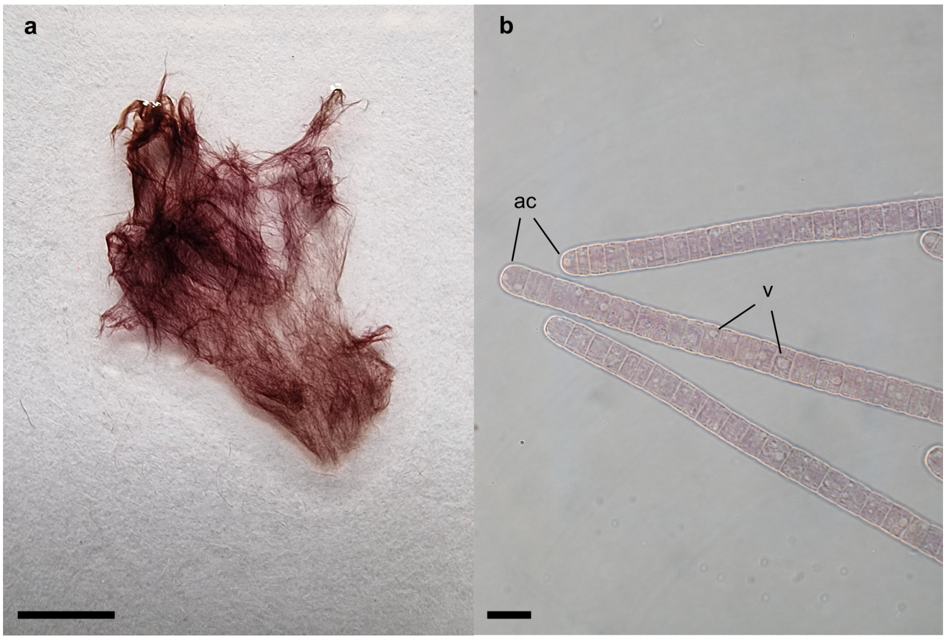


(**a**) Liquid culture of strain MC 67 grown in Z8 medium, showing red-brown biomass, biofilm-like growth on the vessel wall, and cohesive aggregates of filamentous biomass with air pockets adhering to the culture vessel. (**b**) Light microscopy of living culture material showing straight, filamentous trichomes. Cells were arranged in unbranched trichomes without false branching and without visible constrictions at the cross-walls. Apical cells were rounded. Trichomes were 5.5–9.0 $\mu$m wide and individual cells were 1.5–4.5 $\mu$m long. Gliding motility was observed during microscopic examination, whereas akinetes were not detected under the applied cultivation conditions. Designations: ac, apical cells; v, vacuoles. Scale bars: (a) 1 cm; (b) 10 $\mu$m.

### Supplementary Note 1

#### Taxonomic placement of *Microcoleus* sp. MC67.

Phylogenomic inference based on the GTDB bac120 marker set placed MC67 within the *Microcoleus* clade. The closest phylogenetic relative was *Microcoleus bourrellyi* FEM_GT703, and pairwise FastANI analysis yielded an average nucleotide identity of 97.66% between MC67 and FEM_GT703. This value is above the commonly used prokaryotic species-boundary range of approximately 95–96% ANI, supporting a close species-level relationship between both genomes (Jain et al. 2018). The MC67/FEM_GT703 lineage was positioned close to *Microcoleus* sp. BBK16 from Lake Baikal, whereas other anatoxin-associated *Microcoleus* strains were distributed across separate lineages within the broader *Microcoleus* phylogeny.

The close relationship between MC67 and FEM_GT703 is taxonomically relevant because FEM_GT703 was originally described as *Tychonema bourrellyi* from Lake Garda, Italy (Pinto et al. 2017), whereas the GTDB-based framework used here assigns this genome to *Microcoleus* (Salmaso et al. 2016; Shams et al. 2015). This discrepancy highlights the unresolved taxonomic boundary within the *Microcoleus*–*Tychonema* complex. Accordingly, the placement of MC67 should be interpreted within the context of this ongoing taxonomic revision (Skoupý et al. 2026), while the genome-based analysis supports its assignment to the *Microcoleus* lineage used throughout this study.

### Supplementary Note 2

#### Genome assembly, circular replicons and quality assessment

Long-read sequencing enabled the generation of a high-quality genome assembly for strain MC67. The first draft assembly was generated from 3.4 Gbp of filtered Oxford Nanopore reads, with a mean read length of 6,990 bp and a maximum read length of 987 kbp. The final assembly consisted of two contigs: a large 5,609,098 bp chromosome-sized contig with 540× read coverage and a smaller 49,006 bp contig with 108× read coverage. Both contigs were classified as circular by Flye.

Assembly quality assessment supported a near-complete and contamination-free genome. The genome contained 99.1% single-copy BUSCOs, and CheckM2 estimated 98.74% completeness with no detectable contamination. The GC content was 45%, and annotation with Bakta identified 56 tRNAs and 6 rRNAs. Back-mapping of Nanopore reads revealed six high-coverage regions of approximately 2 kb, which were identified as repetitive IS5 insertion sequences.

The use of long-read sequencing was essential for resolving the genome structure of MC67. In particular, the approach enabled circularization of both replicons and allowed the complete anatoxin biosynthetic gene cluster to be assigned unambiguously to the 49 kb circular element rather than to the chromosome-sized contig. This is important because repetitive insertion sequences and secondary replicons can be difficult to resolve using short-read data alone.

### Supplementary Note 3

#### Evidence for a plasmid-like anatoxin biosynthetic element

Several lines of evidence support the interpretation that the 49 kb circular contig represents a plasmid-like or extrachromosomal element. First, the contig was circular and distinct from the 5.61 Mb chromosome-sized replicon. Second, it showed a different read coverage from the chromosome-sized contig, with 108× coverage compared with 508× coverage for the larger replicon. Third, PlasFlow classified the 49 kb sequence as a cyanobacterial plasmid with a probability of 70%. Fourth, the element contained multiple insertion sequence elements flanking the anatoxin biosynthetic genes, consistent with a dynamic genomic environment.

MOB-typer classified the 49 kb element as immobile and did not detect a known origin of transfer or replication type. Therefore, the plasmid-like character of this element cannot be considered fully resolved based on known plasmid markers alone. However, the absence of recognizable replication or mobilization genes may reflect the limited representation of cyanobacterial plasmids, particularly small cyanobacterial plasmids, in current reference databases (Ohdate et al. 2024).

The coverage difference between the 49 kb element and the chromosome-sized contig may indicate that the circular element is maintained at a lower relative copy number than the chromosome. In MC67, the 49 kb element showed approximately 21% of the chromosomal coverage. However, this interpretation requires caution because cyanobacterial genome organization can involve oligo- or polyploidy (Griese et al. 2011; Nies et al. 2020). Filamentous cyanobacteria may contain multiple chromosome copies per cell, and therefore read-coverage differences between chromosome-sized and extrachromosomal elements do not directly translate into absolute copy number without additional experimental validation.

Taken together, circularity, distinct coverage, plasmid-like sequence classification, and the presence of insertion sequences support the interpretation that the anatoxin gene cluster is located on a plasmid-like element. At the same time, the lack of known replication or mobilization genes means that the element is best described conservatively as plasmid-like or extrachromosomal rather than as a fully characterized plasmid.

### Supplementary Note 4

#### Anatoxin gene cluster architecture, toxin profile and evolutionary implications

AntiSMASH identified an anatoxin-like biosynthetic gene cluster on the 49 kb circular element of MC67. The cluster contains the conserved core genes *anaA*–*anaG*, together with the associated genes *anaI*, *anaJ* and *anaK*. This gene content is consistent with previously characterized anatoxin biosynthetic gene clusters, including the canonical cluster of *Oscillatoria* sp. PCC 6506, the dihydroanatoxin-a-producing cluster of *Cylindrospermum stagnale* PCC 7417, and the congeneric cluster of *Microcoleus anatoxicus* PTRS3.

The MC67 cluster showed high similarity to the dihydroanatoxin-a-associated cluster of *Microcoleus anatoxicus* PTRS3. Amino-acid similarities for key genes ranged from 95% to 100%, and both *Microcoleus* clusters contained an SDR family oxidoreductase and a GNAT family N-acetyltransferase downstream of *anaG*. The cluster retained the conserved *anaB–anaG* organization, whereas *anaH* was not identified in either MC67 or PTRS3. This is consistent with previous observations that *anaH*, a transposase-associated gene, is not universally conserved among otherwise complete anatoxin biosynthetic gene clusters (Brown et al. 2016).

Chemical analysis provided functional support for the biosynthetic capacity inferred from the gene cluster. Among the targeted cyanotoxins, only anatoxin-a (ATX) and dihydroanatoxin-a (dhATX) were detected in MC67 cultures. dhATX was the dominant congener, with concentrations ranging from 192 to 2,420 $\mu$g kg$^{-1}$ sample wet mass, whereas ATX occurred at much lower concentrations ranging from 0.22 to 12.16 $\mu$g kg$^{-1}$ sample wet mass. This toxin profile indicates that the pathway in MC67 predominantly produces the reduced dihydro analogue.

The presence of *anaK*, encoding a putative $F_{420}$-dependent oxidoreductase, is consistent with the dominance of dhATX. In *Cylindrospermum stagnale* PCC 7417, AnaK has been proposed to catalyse a reduction step during dhATX formation (Kust et al. 2020), and a homologous *anaK*-containing architecture is also present in the *Microcoleus anatoxicus* PTRS3 cluster (Conklin et al. 2020). However, AnaK remains a candidate enzyme, and direct genetic or biochemical evidence identifying it as the sole reductase responsible for this reaction is still lacking. The low but detectable amount of ATX in MC67 may therefore reflect a minor unreduced fraction of the anatoxin biosynthetic flux under the applied cultivation conditions.

The absence of detectable homoanatoxin-a (HTX) is consistent with the architecture of *anaG* in MC67. In *Oscillatoria* sp. PCC 6506, AnaG contains an extended methyltransferase domain that has been proposed to contribute to HTX formation (Brown et al. 2016). By contrast, *anaG* is shorter in MC67 and in the other non-HTX-producing strains included in the cluster comparison, consistent with the absence of this extended domain and with the observed toxin profile.

The localization of a complete and functional anatoxin biosynthetic gene cluster on a small 49 kb plasmid-like element is notable because cyanobacterial biosynthetic gene clusters have previously been reported mainly on larger plasmids (Popin et al. 2021). The flanking insertion sequences around key components of the cluster suggest a mobile or formerly mobile genomic environment and may indicate past recombination or rearrangement events. This organization supports the broader interpretation that anatoxin biosynthesis is an evolutionarily flexible trait and that plasmid-like elements may contribute to the dissemination or maintenance of toxin biosynthetic capacity in benthic cyanobacteria.

### Supplementary Note 5

#### Phenol-chloroform bead beating high molecular weight DNA extraction

DNA was extracted using a bead-beating step prior to enzyme digestion followed by a phenol-chloroform extraction method to recover high molecular weight DNA. Cells were harvested via filtration of the culture through a 5 µm filter by applying a vacuum. The filter was washed with $ddH_2O$. Cells were scraped off the filter using a spatula, and the filter was cut into small pieces with sterilized scissors. The cell material and filter pieces were resuspended in 500 µL $ddH_2O$. The harvested cells, including the filter pieces, were transferred into a NucleoSpin Soil Extraction Kit bead tube (MACHEREY-NAGEL GmbH & Co. KG, Düren, Germany) containing 4 mm beads. $ddH_2O$ was added to a total volume of 750 µL. Cell disruption was performed through bead beating in a horizontal bead-beating vortex adapter for 7 min.

The sample was then centrifuged for 3 min at 15,000 g, and the supernatant was transferred into a new Eppendorf tube using a cut-off P1000 pipette tip. For enzymatic lysis, 0.04 volumes of lysozyme (50 mg/mL) were added. The sample was mixed by gently inverting the tube 20 times. In this protocol, 30 µL lysozyme were used. The sample was incubated for 60 min at 37 °C and 900 rpm.

For protein digestion, 300 µL SDS (10 %) were added, corresponding to 0.4 volumes SDS. Subsequently, 0.03 volumes of Proteinase K (20 mg/mL) were added, and the sample was mixed by gently inverting the tube 20 times. In this protocol, 22.5 µL Proteinase K were used. The sample was incubated for 2 h at 56 °C and 900 rpm in a heating block. Afterwards, the sample was cooled to room temperature. Cell debris was removed by centrifugation for 5 min at 10,000 g.

The supernatant was carefully transferred into a fresh Eppendorf tube using a cut-off P1000 pipette tip. Samples with a volume above 750 µL were split. One volume of phenol/chloroform/isoamyl alcohol (25:24:1, v/v/v) was added to the sample and mixed thoroughly by gentle inversion. The sample was centrifuged for 3 min at 10,000 g and 4 °C. The upper aqueous phase was carefully transferred into a fresh tube using a cut-off P200 pipette tip, while avoiding disturbance of the interphase.

To remove residual phenol, one volume of chloroform was added to the recovered aqueous phase and mixed thoroughly by gentle inversion. The sample was centrifuged for 3 min at 10,000 g and 4 °C. The upper aqueous phase was carefully transferred into a new tube using a cut-off P200 pipette tip, avoiding both the interphase and the lower phase. If necessary, the chloroform extraction was repeated once to minimize phenol carryover.

For DNA precipitation, 0.1 volumes of 3 M sodium acetate were added to the aqueous phase. Subsequently, 0.7 volumes of 100 % isopropanol were added. The sample was mixed gently by inversion and incubated for 30 min at room temperature. The precipitated DNA was pelleted by centrifugation for 20 min at 10,000 g and 4 °C. The supernatant was discarded, and the DNA pellet was washed with 300 µL 80 % ethanol. The sample was centrifuged for 5 min at 10,000 g and 4 °C. The supernatant was removed, and the pellet was air-dried for 5–10 min.

To further remove contaminants, the DNA precipitation step was repeated. The DNA was resuspended in 15 µL TE buffer and left to dissolve for 10 min at room temperature. DNA concentrations were measured using a Qubit Flex Fluorometer (Thermo Fisher Scientific, Massachusetts, United States). Fragment length was estimated with a TapeStation 4200 (Agilent, California, United States) according to the manufacturer's protocol.

### Supplementary Note 6

#### Bioinformatic tools and parameter settings

Bioinformatic tools were run with the following parameters. Dorado v5.0.0 was used in super-accurate mode. Filtlong v0.3.1 was applied with a minimum mean quality of Q9, a minimum read length of 2,000 bp, and a length weight of 5. Flye v2.9.5 was run with an assembly coverage of 100 and two polishing iterations. NextPolish v1.4.1 was run in 'best' mode with three reruns and automatic genome size estimation. Short-read polishing was performed with BWA mapping and a maximum read depth of 100, while long-read polishing used a minimum read length of 1 kb, a maximum read depth of 100, and the minimap2 preset 'map-ont'. All other tools were used with default settings unless stated otherwise.

### Supplementary Table 1

**Supplementary Table 1 — Genome assemblies used for phylogenetic reconstruction.** Genome assemblies included in the GTDB bac120 marker-gene phylogeny shown in Fig. 2. Toxin status code: first character, presence of the anatoxin biosynthesis gene cassette; second character, chemical detection of anatoxin congeners. Symbols indicate presence or detection (+), absence or non-detection (-), and unavailable or not tested information (?). n.d., collection year not determined or not available.

| Taxon/strain | Origin | Assembly accession | Year | Toxin status | Reference |
|---|---|---|---|---|---|
| *Kamptonema formosum* PCC 6407 | USA, freshwater | GCF_000332155.1 | 1964 | ++ | Shih et al. 2013 |
| *Microcoleus* sp. HI-ES | Iraq, Mosul Dam Lake | GCA_026891855.1 | n.d. | +? | Saeed et al. 2024 |
| *Microcoleus* sp. PH2017_10_PVI_O_A | USA, South Fork Eel River | GCF_020738635.1 | 2017 | ?? | Bouma-Gregson et al. 2021 |
| *Microcoleus* sp. CAWBG51 | New Zealand, Mangatainoka Stream | GCF_020883075.1 | n.d. | – | Tee et al. 2021 |
| *Microcoleus* sp. CAWBG52 | New Zealand, Makarewa River | GCF_020882985.1 | n.d. | – | Tee et al. 2021 |
| *Microcoleus* sp. CAWBG50 | New Zealand, Pelorus River | GCA_020883095.1 | n.d. | – | Tee et al. 2021 |
| *Microcoleus* sp. CAWBG24 | New Zealand, Ashley River | GCF_020883175.1 | n.d. | – | Tee et al. 2021 |
| *Microcoleus* sp. BBK16 | Russia, Lake Baikal | GCF_021648855.1 | 2016 | ?? | Evseev et al. 2023 |
| *Microcoleus bourrellyi* FEM_GT703 | Italy, Lake Garda | GCF_002412335.2 | 2014 | ++ | Pinto et al. 2017; Salmaso et al. 2016 |
| *Microcoleus* sp. MC67 | Germany, Reservoir Mandichosee | This study | 2021 | ++ | This study |
| *Microcoleus* sp. W4-B | Canada, Wolastoq River | GCA_041354795.1 | 2018 | -+ | Valadez-Cano et al. 2023 |
| *Microcoleus* sp. W3-H | Canada, Wolastoq River | GCA_041354945.1 | 2018 | – | Valadez-Cano et al. 2023 |
| *Microcoleus anatoxicus* Micana_A | Switzerland, Areuse River | GCF_042465725.1 | n.d. | ++ | Junier et al. 2024 |
| *Microcoleus* sp. W3-B | Canada, Wolastoq River | GCA_041354715.1 | 2018 | ++ | Valadez-Cano et al. 2023 |
| *Microcoleus* sp. CAWBG640 | New Zealand, Cardrona River | GCA_020883015.1 | n.d. | ++ | Tee et al. 2021 |
| *Microcoleus* sp. EPA2 | USA, East Fork Little Miami River | GCA_020882975.1 | n.d. | – | Tee et al. 2021 |
| *Microcoleus* sp. CAWBG506 | New Zealand, Hutt River | GCA_020883125.1 | n.d. | ++ | Tee et al. 2021 |
| *Microcoleus anatoxicus* PTRS3 | USA, Russian River | GCF_037911295.1 | 2015 | ++ | Conklin et al. 2020 |
| *Microcoleus anatoxicus* PTRS1 | USA, Russian River | GCF_037911335.1 | 2015 | ++ | Conklin et al. 2020 |

### Supplementary Table 2

**Supplementary Table 2 — Reference sequences used for anatoxin biosynthetic gene cluster comparison.** Sequences used for the comparative analysis of anatoxin biosynthetic gene clusters shown in Fig. 1. For each entry, the table lists the strain or organism name, NCBI GenBank assembly or database entry, accession number, nucleotide region used for comparison and reference.

| Name | NCBI GenBank assembly | Accession number | Region [bp] | Reference |
|---|---|---|---|---|
| *Cylindrospermum stagnale* PCC 7417 | GCA_000317535.1 | CP003642.1 | 10829–35497 | Méjean et al. 2016 |
| *Oscillatoria* sp. PCC 6506 | FJ477836 | BGC0000017 | 1–24082 | Méjean et al. 2009 |
| *Microcoleus anatoxicus* PTRS3 | GCA_037911295.1 | JBBLXR000000000 | 1–25072 | Conklin et al. 2020 |
| *Microcoleus* sp. MC67 | This study | This study | 42213–13979 | This study |
